# A capsular polysaccharide-expressing live vaccine suppresses streptococcal toxic shock-like syndrome and provides sequence type-independent protection during *Streptococcus suis* infection

**DOI:** 10.1101/335349

**Authors:** Zhiwei Li, Peixi Chang, Jiali Xu, Chen Tan, Xiaohong Wang, Weicheng Bei, Jinquan Li

**Affiliations:** Bio-Medical Center, Key Laboratory of Environment Correlative Dietology, College of Food Science and Technology, Huazhong Agricultural University, Wuhan, Hubei, People’s Republic of China; State Key Laboratory of Agricultural Microbiology, Huazhong Agricultural University, Wuhan, Hubei, People’s Republic of China

## Abstract

*Streptococcus suis* (*S. suis*) is an encapsulated zoonotic pathogen, which is responsible for bacterial meningitis and streptococcal toxic shock-like syndrome (STSLS). Despite many attempts to develop an effective vaccine, none is currently available. Here, a capsular polysaccharide (CPS)-expressing attenuated mutant 2015033 was constructed by deleting five virulence-associated factors (*sly*, *scpA*, *ssnA*, *fhb*, and *ssads*) in an outbreak *S. suis* strain SC19. Genes mentioned above are associated with either innate immunity-evading or tissue barrier-invading. Deletion of these genes did not impact the growth ability and CPS generation of 2015033, and the mutant exhibited no hemolytic activity to erythrocytes and no cytotoxicity to different epithelial or endothelial cells. In addition, 2015033 was more easily eliminated by whole human blood *in vitro* and by mouse blood *in vivo*. In addition, 2015033 showed a diminished invasive ability in different mouse organs (brain, lung, and liver) and avirulent properties in mice associated with weak inflammation-inducing ability. Immunization with 2015033 triggered T cell-dependent immunity and this immunity suppressed STSLS during SC19 infection by inhibiting excessive proinflammatory responses. In addition, immunization with 2015033 successfully conferred sequence type (STs)-independent protection to mice during heterogeneous infections (ST1, ST7, and ST658). This study presents the feasibility of the strategy of multi-gene deletion for the development of promising live vaccines against invasive encapsulated pathogens.

**IMPORTANCE:** *S. suis* is a traditional zoonotic agent causing human meningitis and STSLS, which is also a neglected emerging food-borne pathogen. Increasing antimicrobial resistance invokes reduction of preventative use of antibiotics in livestock creating an urgent need for effective vaccines. Given the expression of CPS is the basis for promising vaccines against encapsulated pathogens, and in order to find an effective and economical strategy for CPS-based vaccine development, multi-gene deletion was introduced into the design of a *S. suis* vaccine for the first time. From our results, CPS-expressing attenuated mutant 2015033 exhibited diminished evasive ability against the innate immune system and reduced invasive properties against different host barriers. To our knowledge, 2015033 is the first STSLS-suppressing *S. suis* vaccine to provide STs-independent protection during heterogeneous infections.

## INTRODUCTION

*Streptococcus suis* (*S. suis*) is an important zoonotic pathogen with considerably high phenotypic heterogeneity, which can be classified into 33 serotypes based on the antigenicity of their capsular polysaccharides (CPS) (1) and 1003 sequence types (STs) according to the sequence of seven house-keeping genes (*cpn60*, *dpr*, *recA*, *aroA*, *thrA*, *gki*, and *mutS*) (2). As the most virulent serotype, *S. suis* serotype 2 (*S. suis* 2) has caused a wide variety of diseases in pigs, including sepsis, meningitis, and arthritis, leading to economic losses in the porcine industry. Moreover, this pathogen is also an emerging agent of severe human infection cases around the world, causing meningitis and streptococcal toxic shock-like syndrome (STSLS) (3, 4). The first *S. suis* infection case of human was reported in Denmark in 1968, and to date, there are more than 1600 described human infection cases including two large outbreaks (1998 and 2005) in China (5). Antimicrobials should have been an effective method to prevent *S. suis* infection; however, a reduction in preventative antibiotics in livestock worldwide has increased the need for *S. suis* vaccines (6).

Because of the limited success of commercial bacterins (7, 8), many new strategies including subunit and live vaccines have been experimentally evaluated. However, the application of subunit vaccines is restrained as these vaccines are dependent on adjuvant and only provide partial or no cross-protection against *S. suis* strains with different STs (9). Hence, several attempts were made to explore adjuvant independent live vaccines against *S. suis*, which also revealed potential to confer host cross-protection against heterologous *S. suis* strains (10). In addition, given the many *in vivo*-induced genes of *S. suis* that are usually upregulated or only expressed during infection (11), live vaccines possess incomparable advantages to trigger specific immunity against these *in vivo*-induced factors.

Encapsulated pathogens like *S. suis* employ CPS to evade host innate immunity (12). CPS is the first exposed layer to the host and is associated with functions of various bacterial components and with the immunogenicity of vaccine strains (13), implying the importance of CPS expression for vaccines against encapsulated pathogens. In fact, it has been reported that once anti-CPS antibodies are produced, they will elicit a strong potential to protect the host from *S. suis* challenge (14, 15).

As T-cell independent antigens, the poor immunogenicity of carbohydrates is one of the main obstacles hindering the development of CPS-based vaccines (13, 16). It is well known that protein-carbohydrate conjugation is an important strategy to induce long-lasting protection against encapsulated bacteria (16, 17). However, glycoconjugate vaccines may need to be polymerized or modified for effective protection and are usually complex and expensive (16, 18). Hence, applying genetically modified attenuated strains that simultaneously express CPS and various other proteins (a major class of molecular targets for antibody therapies) (13) is a promising and cost-effective strategy for the development of vaccines against encapsulated pathogens.

However, the absence of vaccine candidate strains, which are fully attenuated but can still induce potent protective responses limits the application of live vaccines against *S. suis* (10). Pathogenesis of *S. suis* is complicated that there are more than 100 virulence-associated factors have been reported, but none of them is considered critical virulent factor (19, 20), making it difficult to obtain an ideal vaccine candidate by single-gene deletion. Indeed, except CPS deficient mutants, other gene deletion mutants have been proven to be insufficiently attenuated (21-23), while CPS deficient strains conferred only minor protection (24).

Considering that evading innate immunity and invading tissue barriers are the two most important processes during *S. suis* infection, with the premise of maintaining CPS, we selected four genes including *scpA*, *ssnA*, *fhb*, and *ssads* to construct a live vaccine strain. These four genes are involved in the evasive properties of the pathogen against host innate immune responses (25-29), which are responsible for STSLS induced by the streptococcal inflammatory reaction (30). Among these genes, *scpA* and *fhb* help the pathogen elude complement-mediated immune defenses (27, 28), while *ssnA* and *ssads* are inhibitors of neutrophil-mediated immune responses (25, 29). In addition, deletion of *sly*, a suilysin encoding gene was also taken into account for live vaccine strain construction; SLY is the only well-characterized *S. suis* toxin and induces rupture of erythrocytes leading to the release of cellular contents (31); s*ly* was also shown to promote migration of *S. suis* through the blood brain barrier (32).

In this study, all five genes mentioned above were deleted in an outbreak *S. suis* 2 strain, SC19 (33), to obtain an avirulent, CPS-expressing vaccine candidate. This five-gene deletion mutant was renamed as 2015033 and the safety of 2015033 was determined both *in vitro* and *in vivo*. Furthermore, the immune effects of 2015033 on the prevention of STSLS and acute death during heterogeneous infections were also evaluated in mice, illuminating the feasibility of multi-gene deletion strategies in the development of vaccines against encapsulated pathogens.

## RESULTS

### Growth ability and CPS expression of 2015033 were unaffected by gene deletion

To generate a fully attenuated live vaccine strain, five virulence-associated genes that play different but important roles were deleted in an outbreak *S. suis* 2 strain, SC19 (Fig. 1); the subsequent mutant (Δ*sly*Δ*scpA*Δ*ssnA*Δ*fhb*Δ*ssads*) was renamed as 2015033. The successful gene-deletion was confirmed by PCR and further verified by sequencing (Fig. S1). As a low level replication of a vaccine strain obviously limited its effect (10), it was necessary to maintain normal growth ability in 2015033. The growth curve of 2015033 was found to be relative to SC19 during incubation, and the bacterial count of 2015033 was also similar to SC19 at the stationary phase (Fig. 2A). Previously, a nonencapsulated *S. suis* strain induced only vulnerable immunity (24), while our CPS generation of 2015033 was not impaired by gene deletions (Fig. 2B). In addition, the MIC of penicillin against 2015033 was the same as that for SC19 (0.16 μg/mL), implying cell wall phenotype of 2015033 was unaffected (Fig. 2C). Taken together, these results demonstrated that deletion of five genes had no negative impact on the basic biological characteristics of 2015033, which guarantee the efficiency of the vaccine strain.

**FIG 1.**
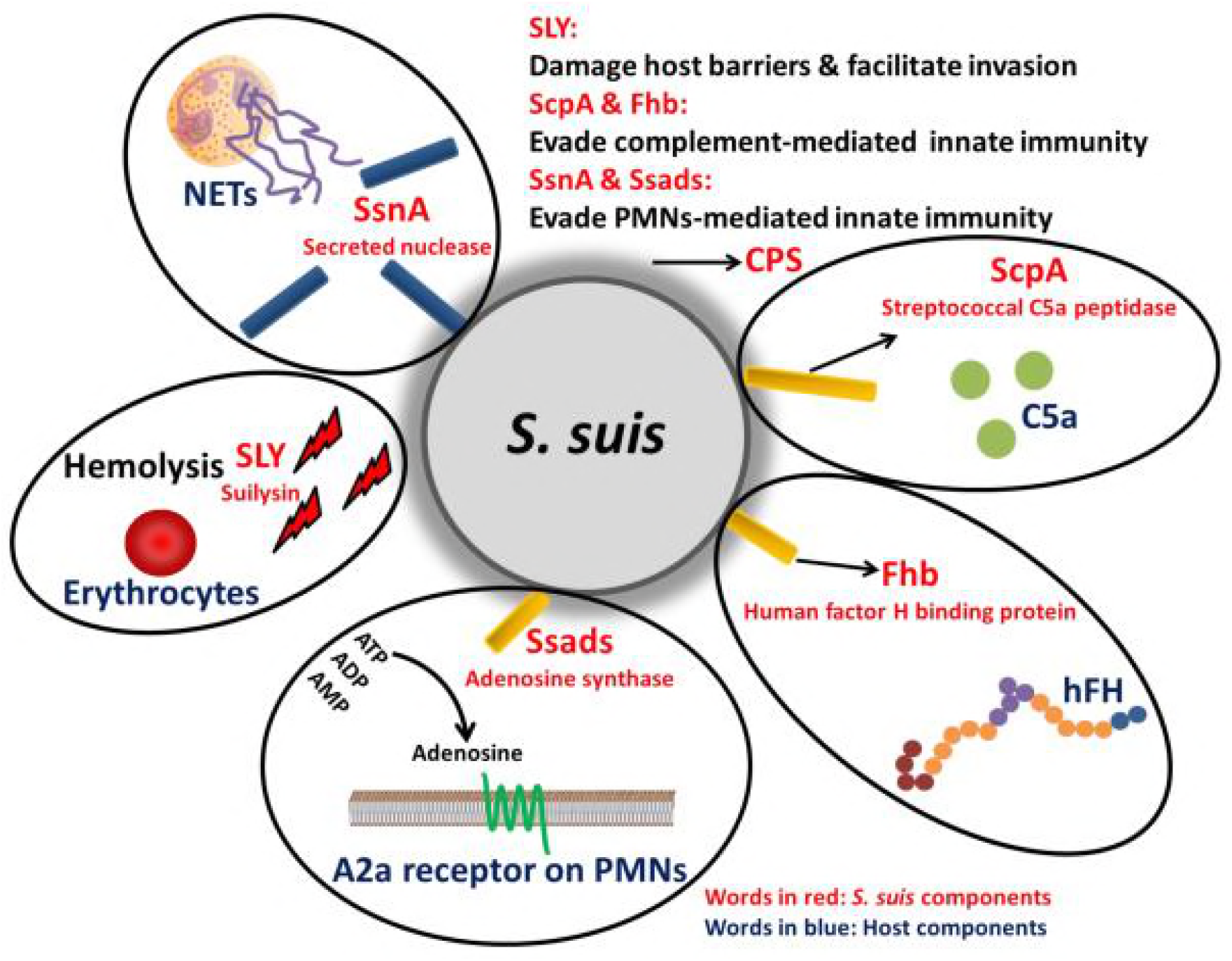
The five genes deleted in this study were associated with abilities to cross host tissue barriers and to evade host innate immune defenses. SLY (SSUSC84_1264) is a thiol activated cytolysin, which promotes invasion by forming pores in eukaryotic cell membranes, resulting in the release of cellular contents. ScpA (SSUSC84_1795) is a surface associated subtilisin like serine protease, which is homologous to the streptococcal C5a peptidase of GAS, which inactivates chemotactic factor C5a and C5a-mediated innate immunity. Fhb (SSUSC84_0242) is a human factor H (hFH) binding protein which interacts with human complement regulator hFH to inhibit complement activation and complement-mediated phagocytosis. SsnA (SSUSC84_1782) is the first reported specific neutrophil extracellular traps (NETs)-evading component of *S. suis*. Ssads (SSUSC84_0906) is an adenosine synthase, which impairs polymorphonuclear leukocytes (PMNs)-mediated innate immunity by reducing the oxidative activity and the degranulation of PMNs via A2a receptor. *S. suis* equipped with these five virulence-associated factors can be described as a “small tank”, which can simultaneously evade innate immune defenses and attack host barriers to facilitate its invasion and survival.

**FIG 2.**
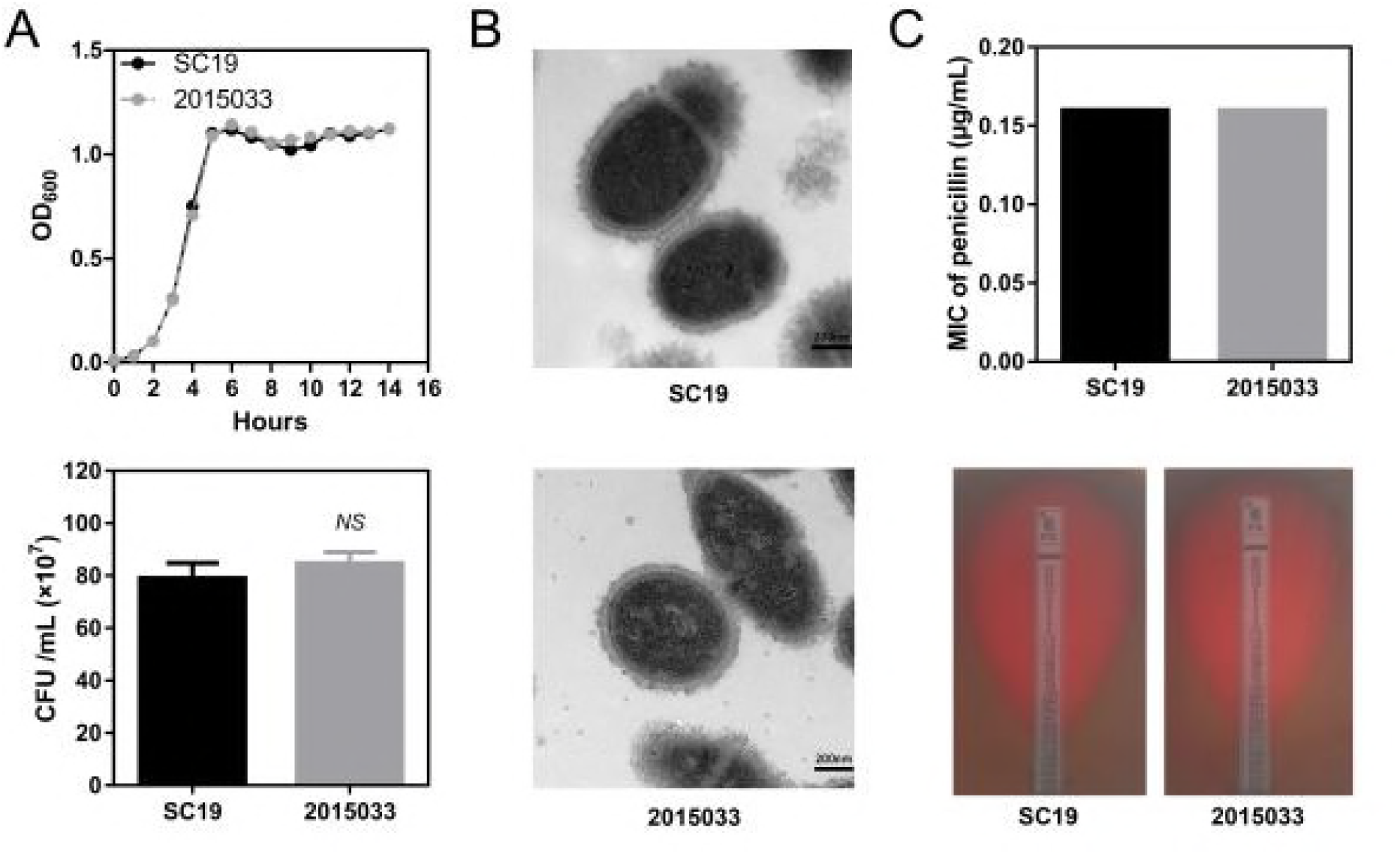
Basic biological characteristics of 2015033. (A) Growth ability: bacteria were incubated at 37 °C and the optical density (600 nm) of the culture was measured at a 1-h interval. At 6 h post inoculation, the bacterial counts in cultures were calculated. (B) Transmission electron micrographs. The bar indicates the magnification size 200 nm. (C) Penicillin *E*-test on Mueller Hinton agar. *E*-test strips were used with a penicillin concentration gradient in the range of 0.02 μg/mL to 32 μg/mL. *NS*, No Significance.

### Evasive and invasive abilities of 2015033 were remarkably attenuated *in vitro*

*S. suis* hired various strategies to evade innate immune defense and proliferate rapidly in the host, inducing excessive inflammatory responses (29). Whole blood killing assay showed the growth rate of 2015033 was much lower than that of SC19 in human blood, indicating the immune escape ability of the mutant was significantly decreased (Fig. 3A). As expected, SLY deficient 2015033 showed little hemolysis on sheep erythrocytes while SC19 induced obvious hemolytic effects (Fig. 3B). SC19 induced obvious cytotoxicity to human brain microvascular endothelial cells (HBMEC), human lung carcinoma cells (A549), and human colorectal adenocarcinoma cells (Caco-2), while 2015033 showed no obvious effects (Fig. 3C). As *S. suis* is a neuro-invasive pathogen, which is known to promote BMEC injury initiating entry into the central nervous system (34), the interaction between 2015033 and HBMEC was further explored. Cells treated with SC19 showed significant necrotic features (PI^+^/Annexin V-FITC^+^), while almost all cells treated with 2015033 were normal (PI^-^/Annexin V-FITC^-^) (Fig. 3D). These results indicated the invasive ability that facilitates the migration of *S. suis* across different barriers was dramatically reduced in 2015033. Although 2015033 showed no cytotoxicity, its adhesive capacity to HBMEC was similar to that of SC19 (Fig. 3E), implying adhesins in 2015033 were well preserved.

**FIG 3.**
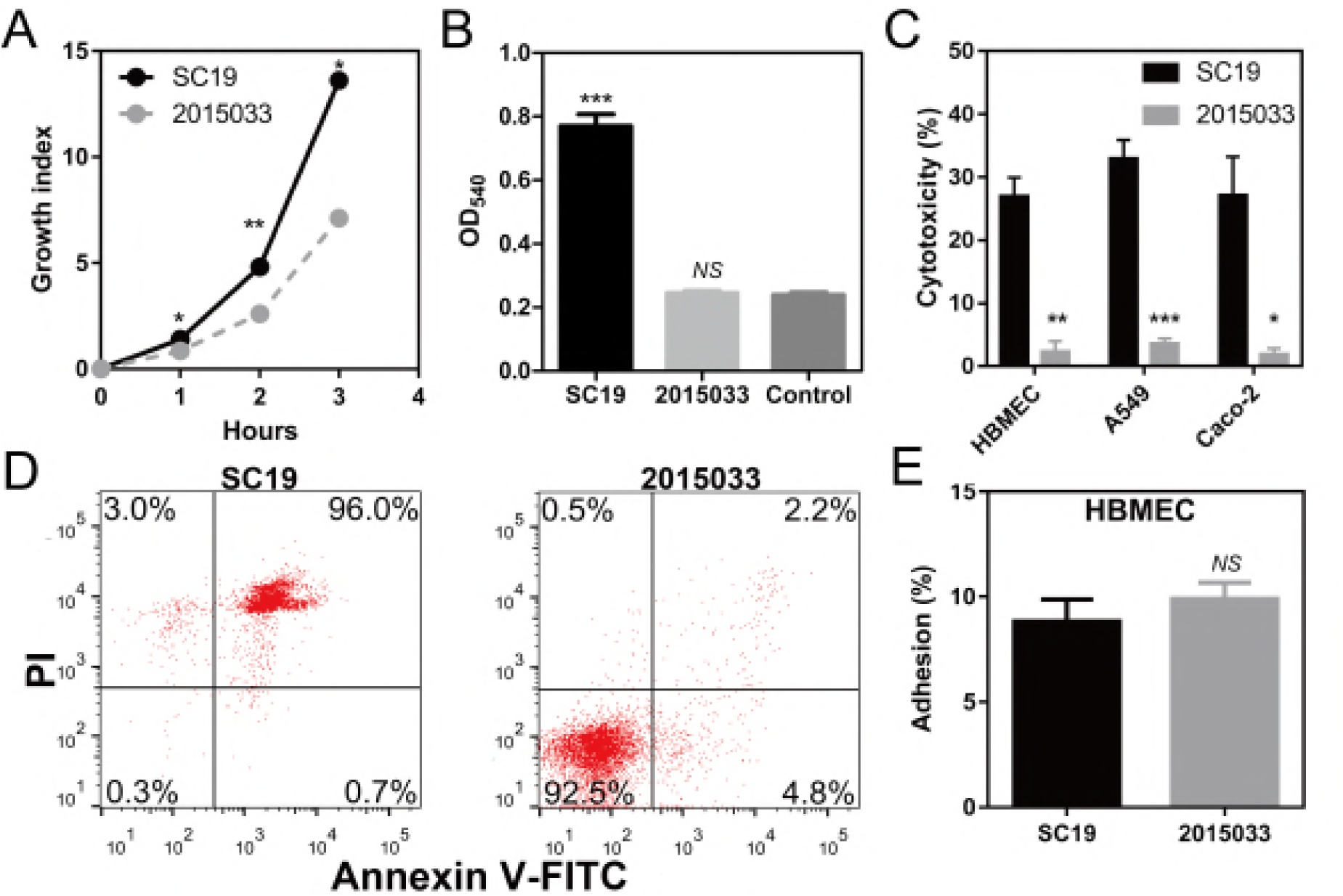
*In vitro* virulence test of 2015033. (A) The growth of 2015033 in human blood: diluted cultures of bacteria (100 μL) were added into whole human blood (900 μL) for incubation. Samples were plated on TSA to determine bacterial counts. (B) Hemolytic activity of 2015033: 100 μL cultures of bacteria or sterile culture were separately incubated with 900 μL 4% sheep erythrocytes for 1 h at 37 °C. The mixtures were then centrifuged and the optical density (540 nm) of each supernatant was measured. (C) Cytotoxicity of 2015033 to different barrier cells: bacteria were incubated with different cells (MOI = 20) for 4 h at 37 °C. LDH measurements were performed using an LDH-cytotoxicity assay kit. (D) Cell death model of HBMEC after *S. suis* infection: bacteria were incubated with HBMEC (MOI = 20) for 8 h at 37 °C, PI/Annexin V-FITC double-staining was used to determine subsequent cell death; the percentage cells in each region were then labeled. (E) Adherence to HBMEC by 2015033: bacteria were incubated with HBMEC (MOI = 10) for 3 h at 37 °C and the cells were lysed with 1% saponin. Lysates were then plated on TSA to determine the bacterial adhesion. *NS*, No Significance; \**p* < 0.05; \*\**p* < 0.01; \*\*\**p* < 0.001.

### 2015033 exhibited avirulent capabilities *in vivo* and elicited weak inflammation-inducing ability

Having confirmed that 2015033 was remarkably attenuated *in vitro*, we then used BALB/c mice to evaluate the pathogenicity of 2015033 *in vivo*. All mice infected with SC19 died within 2 days, in contrast to a survival rate of 100% in the 2015033 challenged mice (Fig. 4A). To explore the mechanisms driving increased survival post 2015033 infection, *in vivo* colonization experiments were carried out. 2015033 counts isolated from the brain at 24 h post infection and from blood, lung, brain, and liver at 72 h post infection were significantly lower than that of SC19 (Fig. 4B). As mice challenged with SC19 usually died approximately 18 hours post-infection (Fig. 4A), we used lower infectious doses to determine whether inflammatory responses were associated with these quick mortality rates. The levels of six inflammation-associated cytokines (IL-6, IL-10, IL-12p70, IFN-γ, TNF-α, and MCP-1) were measured at 18 hours post-infection, and production of the above cytokines in 2015033 infected mice was much lower than SC19 infected mice (Fig. 4C), which helped to explain the substantial attenuation of 2015033. In summary, 2015033 exhibited avirulent features *in vivo* accompanied with substantially subdued inflammation-inducing abilities, which further revealed the safety of 2015033 for use as a vaccine.

**FIG 4.**
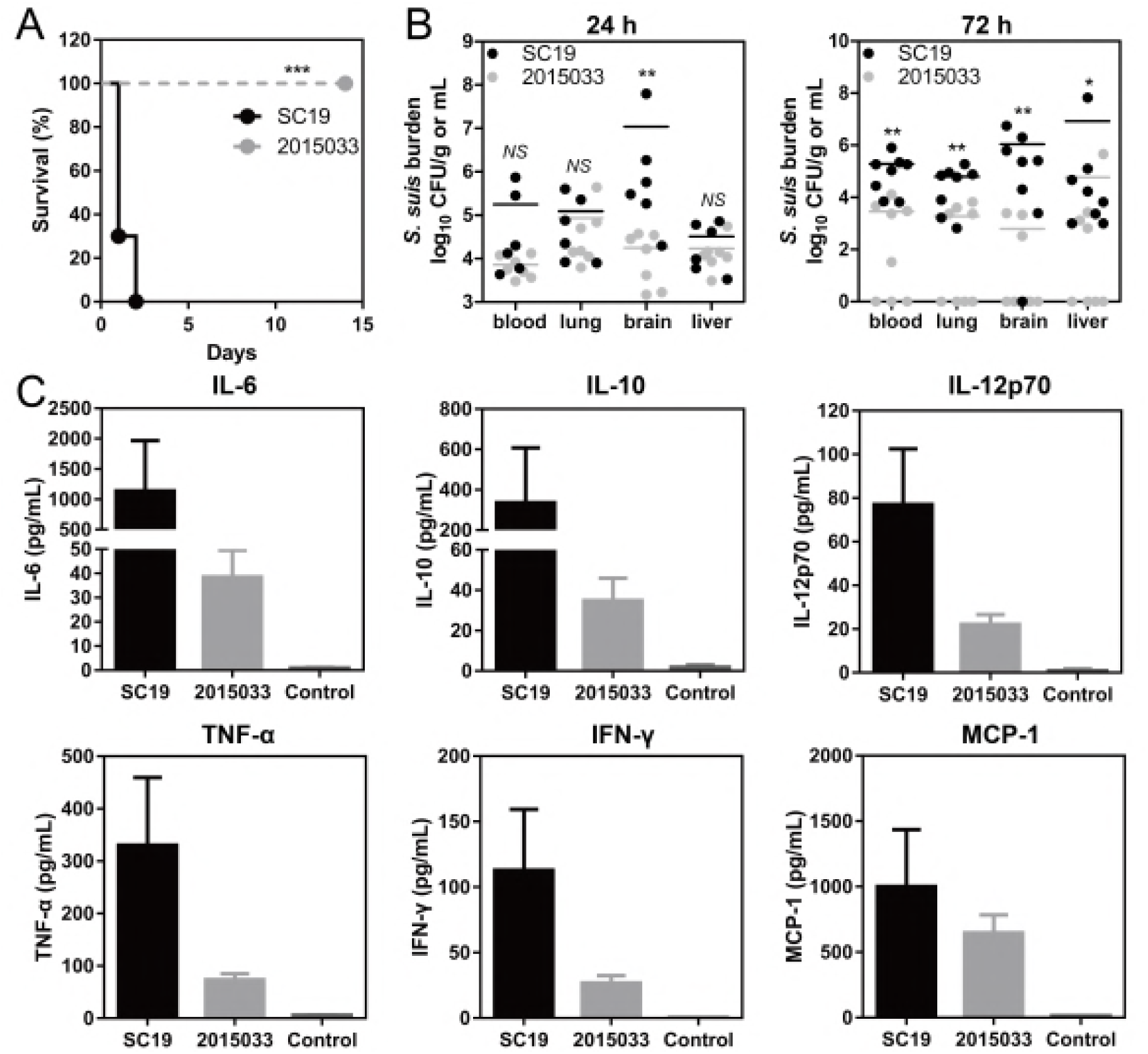
*In vivo* virulence test of 2015033. (A) Survival of mice infected with SC19 or 2015033: mice (six-week-old female, ten mice per group) were intraperitoneally infected with SC19 or 2015033 (2 × 10^8^ CFU) and were monitored daily up to 14 days post-infection. (B) Bacterial burdens in blood, lung, brain, and liver of mice: mice were intraperitoneally infected with SC19 or 2015033 (5 × 10^7^ CFU) and their tissues were isolated at 24 h (seven mice) and at 72 h (eight mice) post-infection to determine bacterial burdens. Results from individual mice are shown as log 10 of bacterial counts (CFU/g or CFU/mL). (C) Cytokine assessment: mice were intraperitoneally infected with SC19 or 2015033 (5 × 10^7^ CFU) and serum samples were collected by cardiac puncture at 18 h post infection. Levels of IL-6, IL-10, IL-12p70, IFN-γ, TNF-α, and MCP-1 were assessed using the CBA mouse inflammation kit. *NS*, No Significance; \**p* < 0.05; \*\**p* < 0.01; \*\*\**p* < 0.001.

### Immunization with 2015033 induced T cell-dependent responses to *S. suis*

As isotype-switched IgG plays an important role in the fight against encapsulated extracellular bacteria (35, 36), antibodies (Abs) to the entire encapsulated SC19 strain were assessed after 2015033 immunization. Immunized mice exhibited a significant increase in IgG, IgG1, and IgG2a, which remained at high levels until the end of the experiment, while mice injected with phosphate buffered saline (PBS) showed no detectable increases (Fig. 5A). Meanwhile, there was a significant increase in IgG, IgG1, and IgG2a specific to an isogenic nonencapsulated mutant Δ*cps2E* (Fig. 5B). Significant Ab titers specific to Δ*cps2E* were also observed post immunization and remained at high levels throughout the experiment (Fig. 5C). In addition, the IgG1 response was significantly higher than that of IgG2a regardless of the antigen type, suggesting Th2-type bias (Fig. 5). Abs detection demonstrated that immunization with 2015033 induced a two-pronged T cell-dependent immune reaction specific to both *S. suis* 2 CPS and proteins.

**FIG 5.**
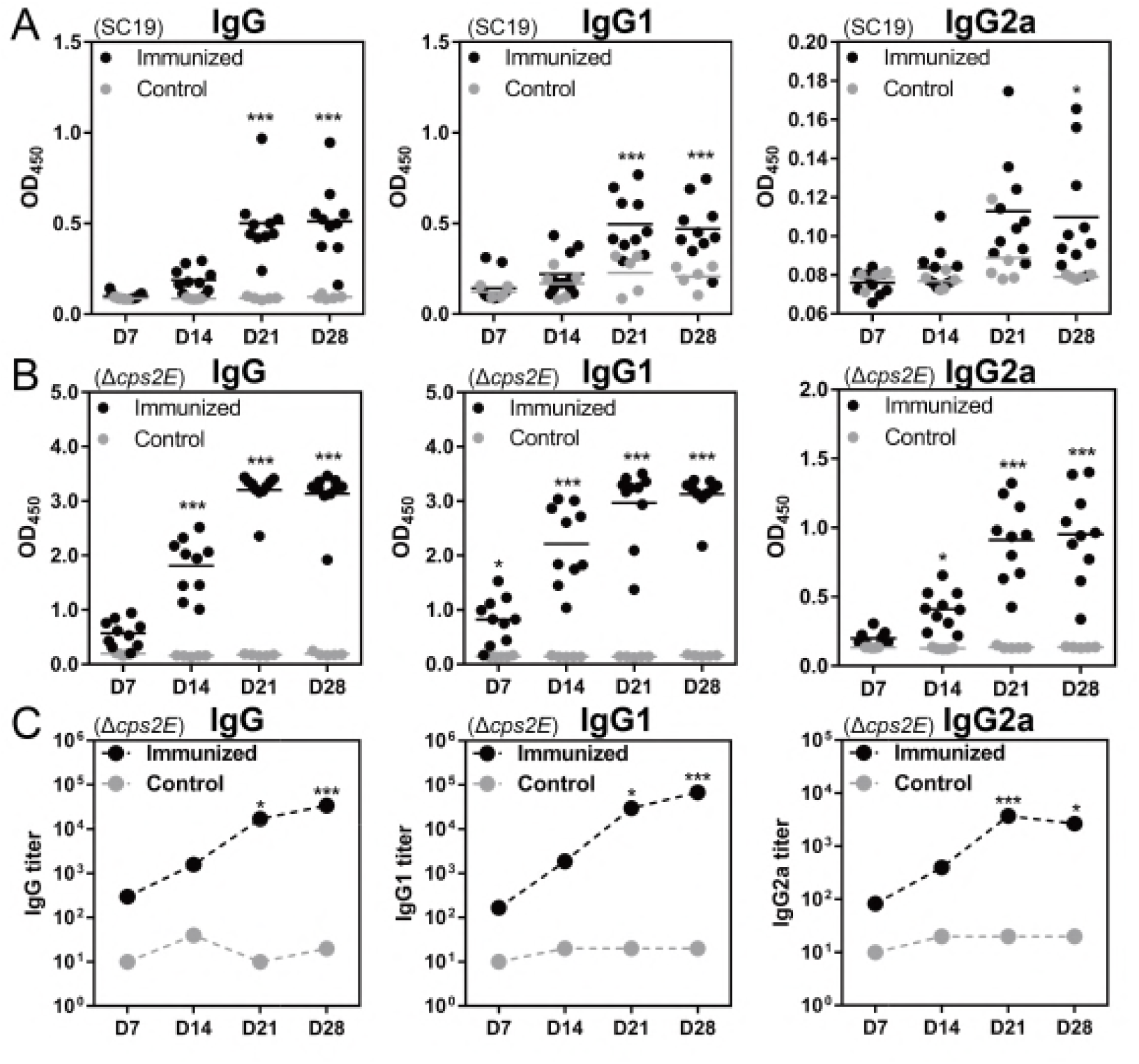
Ab responses after 2015033 immunization. Mice (four-week-old female) were subcutaneously immunized twice with 2015033 (2 × 10^7^ CFU, suspended in 200 μL PBS) at a 2-week interval, while negative controls were injected with PBS. On days 7, 14, 21, and 28 serum samples were taken for ELISA. (A) Quantification of SC19-specific Ab responses: after pre-coated microtiter plates with SC19, serum samples at the respective time points were plated at a dilution of 1/50 in 5% skim milk and the absorbance was read at 450 nm. (B) Quantification of protein-specific Ab responses: microtiter plates were pre-coated with nonencapsulated mutant Δ*cps*2E developed from SC19 and processed directly as above (panel A). (C) Titration of protein-specific Abs: mouse sera were serially diluted (2-fold) in 5% skim milk and processed as above (panel A). Significant differences between immunized and control groups were defined as \**p* < 0.05 and \*\*\**p* < 0.001.

### Immunization with 2015033 suppressed STSLS induced by the outbreak strain

Excessive in?ammatory responses caused by *S. suis* are a hallmark of STSLS (30); therefore, in order to gain insight into the potential role of 2015033 immunization in suppressing *S. suis*-induced STSLS, the levels of six inflammation-associated cytokines (IL-6, IL-10, IL-12p70, IFN-γ, TNF-α, and MCP-1) were measured (Fig. 6A). After challenge with the outbreak strain SC19, all six inflammatory cytokines were obviously increased in PBS treated mice, but only slight increases were observed in 2015033 immunized mice (Fig. 6B), indicating immunization with 2015033 successfully suppressed inflammatory responses during infection.

**FIG 6.**
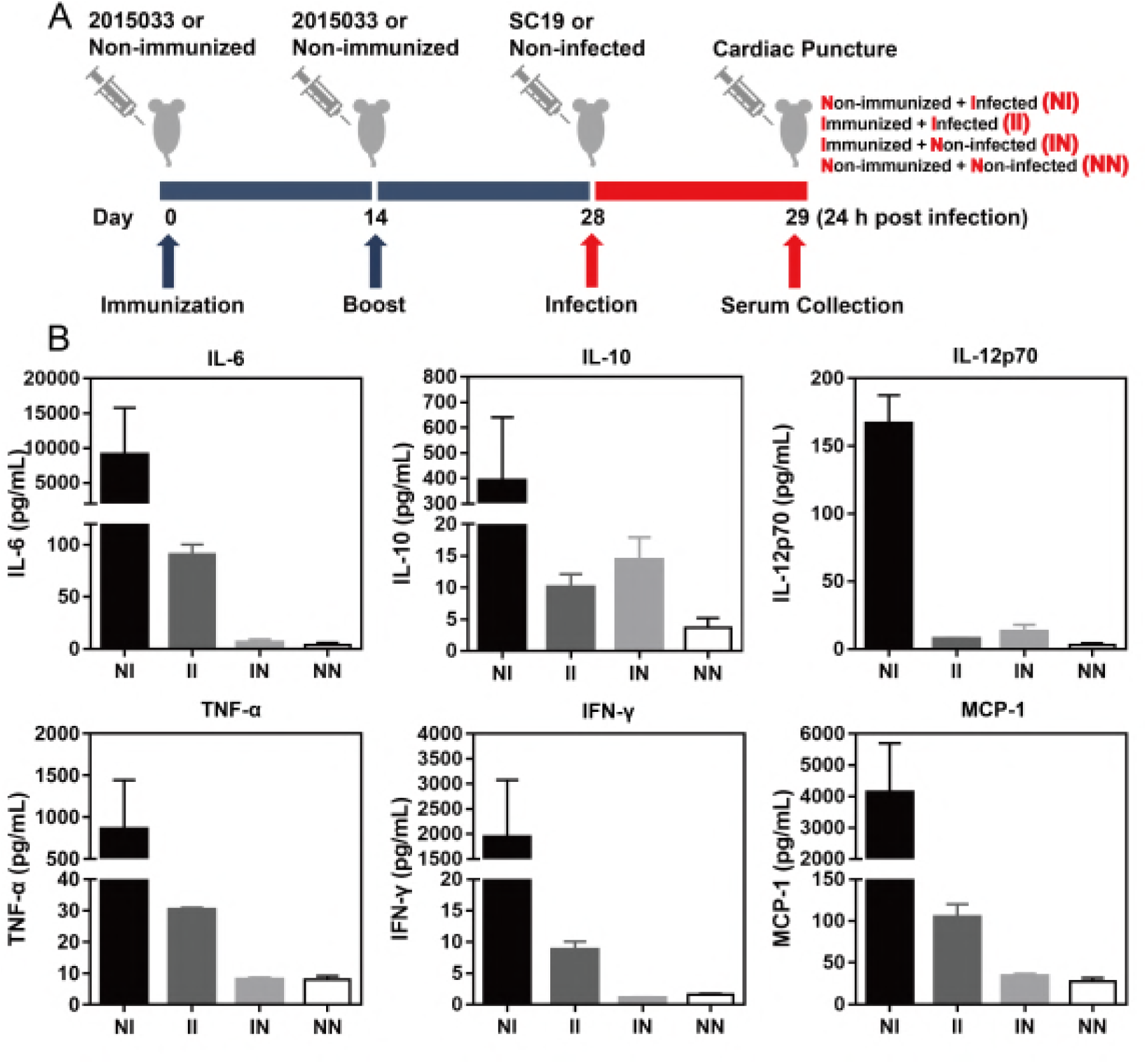
Inflammatory responses of immunized mice after SC19 challenge. (A) Schematic design of this study: mice (four-week-old female) were subcutaneously immunized twice with 2015033 (2 × 10^7^ CFU, suspended in 200 μL PBS) at a 2-week interval, while non-immunized mice were injected with PBS. On day 28, mice were intraperitoneally infected with SC19 (5 × 10^7^ CFU, suspended in 200 μL PBS) or PBS. At 24 h post-infection serum samples were collected by cardiac puncture for cytokine detection. (B) Cytokine detection according to the scheme shown in panel A. Levels of IL-6, IL-10, IL-12p70, IFN-γ, TNF-α, and MCP-1 were assessed using CBA mouse inflammation kit. Results are representative of two independent experiments.

### Immunization with 2015033 protected mice from heterogeneous infections

STs belonging to the ST1 clonal complex (CC1) are primarily associated with disease in both animals and humans around the world (Fig. 7A). We employed three strains (SC19, P1/7, and LSM102) with different STs (ST7, ST1, and ST658) to evaluate the cross-protection conferred by 2015033 (Fig. 7A). After immunization, the opsonic killing assay was conducted. We found that growth of SC19, P1/7, and LSM102 were significantly inhibited by pretreatment with immune serum (Fig. 7B). Virulent challenge showed all mice that were administrated PBS died within 3 days post SC19 infection, while 90% of the mice immunized with 2015033 survived throughout the experiment (Fig. 7C). Mice in the 2015033 immunized group survived throughout the experiment post P1/7 infection, while survival rate was only 30% in the PBS administrated group (Fig. 7C). In addition, statistically significant protection was also observed after LSM102 challenge; the survival rates of mice were 20% and 70% in the PBS group and 2015033 immunized group, respectively (Fig. 7C). These results showed immunization with 2015033 conferred cross-STs protection in mice against not only an isogenic ST7 strain but also heterogenic ST1 and ST658 stains.

**FIG 7.**
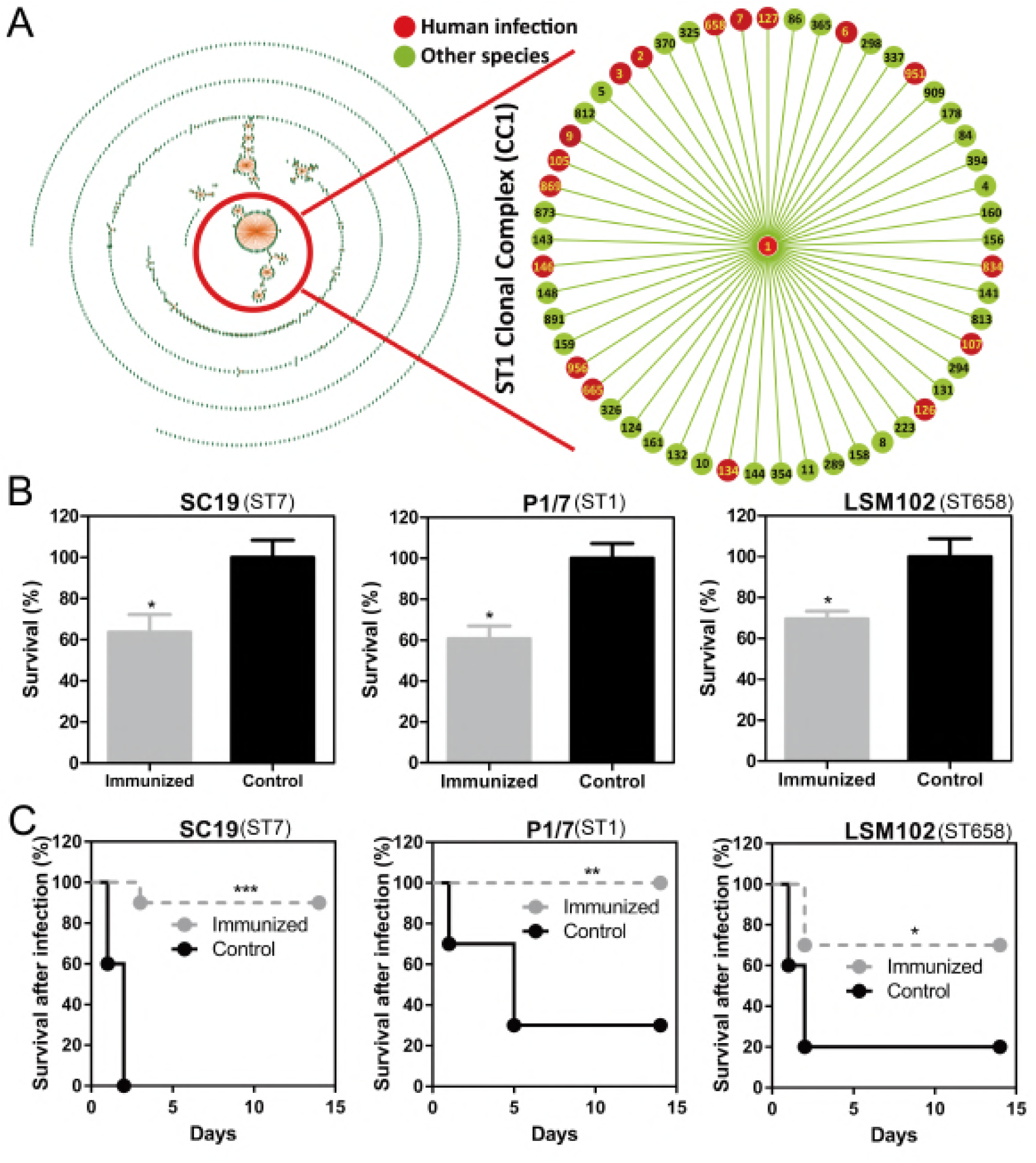
STs-independent protection conferred by 2015033. (A) Population snapshot of 1003 STs of *S. suis*. Two different STs sharing six of the seven housekeeping genes form a single-locus variant (SLV). At least three STs with only one SLV form a clonal complex. Population snapshots of clonal complexes were drawn using eBURST software. STs involved in human infection were labeled as red. (B) The opsonic killing of ST7 strain SC19, ST1 strain P1/7, and ST658 strain LSM102 (shown in panel A). Bacteria (1 × 10^3^ CFU) were incubated in 100 μL immune or nonimmune mouse sera for 30 min at 37 °C. The mixtures then added into 400 μL fresh nonimmune mouse whole blood for 1 h incubation and the diluted samples were plated on TSA to determine bacterial counts. (C) Survival of immunized mice post heterogeneous infections: mice in different groups were intraperitoneally infected with ST7 strain SC19, ST1 strain P1/7, or ST658 strain LSM102 (2 × 10^8^ CFU, suspended in 200 μL PBS, ten mice per group) on day 28 and were monitored daily up to 14 days post infection. Statistical significance for survival study was assessed by log-rank test. \**p* < 0.05; \*\**p* < 0.01; \*\*\**p* < 0.001.

### DISCUSSION

CPS is one of the most important antigens of encapsulated pathogens (16) and since carbohydrates are considered T cell-independent antigens, which mainly elicit IgM without memorability and isotype-switched IgG (16, 18). In this study, we attempted to construct an attenuated immunogenic strain, which simultaneously expressed CPS and various protein antigens, to improve current *S. suis* vaccine development.

Identifying ideal target genes is the first and the most important step in the multi-gene deletion process. Based on previous studies, we supposed that two categories of virulence-associated factors should be given priority when designing *S. suis* live vaccine, one is mainly associated with avoiding immune defense and another is mainly for invading tissue barriers (3, 6). The effective construction of live vaccines should involve reducing the virulence of *S. suis* based on these two aspects and retaining sufficient immunogenicity. Among the five genes deleted in this study, *sly* is considered a cytotoxicity-inducing protein, which aides the pathogen in crossing tissue barriers and which is involved in upregulated inflammatory responses (31, 32). The other four genes, *scpA*, *ssnA*, *fhb,* and *ssads*, were demonstrated to be involved in evading host innate immune defenses (25, 27-29).

Compared with the parental SC19 strain, growth ability and CPS generation of the five-gene deletion mutant 2015033 were not affected, which was important for the vaccine strain to reach an adequate level *in vivo* for the induction of immune responses (10). Furthermore, 2015033 exhibited weak resistance in whole blood killing assay and showed nontoxicity to different tissue barrier cells. Interestingly, 2015033 showed a similar adhesive capacity to HBMECs compared with SC19. This may be due to many important *S. suis* adhesins were well preserved in 2015033 (37, 38). In fact, adhesin-specific Abs are effective in conferring protection against *Streptococcus pyogenes* by blocking the binding ability of the adhesin to host tissue (neutralization) or by tagging the bacteria for phagocytosis (opsonization) (39). As *S. suis* 2 which cannot invade endothelial cells usually secretes SLY to damage the blood brain barrier to facilitate its infection (32), our results implied 2015033, which could not destroy but still maintained the comparable adhesive capacity to host cells, seemed to be an adequate stimulant of immune cells (40). Moreover, because of the deletion of *sly*, 2015033 could not induce injury to immune cells, and therefore, antigen-presenting cells were still able to recognize the bacteria for the activation of further immune responses. These effects not only promoted host immune-inducing abilities but also indicated the safety of the vaccine strain.

Immunization with 2015033 triggered significant isotype-switched IgG. Considering the inability of carbohydrates to induce T cell-dependent responses (13), significant IgG switching in response to SC19 may be explained by the two-pronged (anti-CPS and anti-protein) immunity conferred by 2015033. In the immunized group, four cytokines (IL-6, IFN-γ, TNF-α, and MCP-1) were increased after SC19 challenge, while other two cytokines (IL-10 and IL-12p70) were not. Given that IL-10 is known to be an anti-inflammatory cytokine, it was reasonable that IL-10 was not increased during *S.suis* infection (41). IL-12p70 is known to promote the maturation of naive T cells to helper Th1 cells (42); the unchanged levels of IL-12p70 may support the Th2-type bias after immunization. Inflammation-associated cytokines play important roles in the elimination of bacterial pathogens; yet during *S. suis* infection, inflammation seems to be a two-edged sword as excessive inflammatory responses are also responsible for STSLS. We found 2015033 immunized mice had slight increases in levels of inflammatory cytokines after SC19 challenge, and it needs to be determined if this inflammatory response is essential for the host to eliminate *S. suis*. If this proves to be the case, it will also be necessary to determine the boundary between the protective inflammatory response and fatal inflammatory response during *S. suis* infection.

Based on the sequence of seven house-keeping genes (*cpn60*, *dpr*, *recA*, *aroA*, *thrA*, *gki*, and *mutS*), *S. suis* could be classified into 1003 STs. Although a serotype 9 strain has been demonstrated to give cross-protection against different STs belonging to CC16 (43), there is no live vaccine reported that provides cross-protection against different STs in CC1, which is considered the most virulent and extensive clonal complex (5). In this study, three STs belonging to CC1 were used: ST1 predominates in many European and Asian countries while ST7 is considered to be responsible for two outbreaks in China (5). ST658 (LSM102) is a newfound virulent ST isolated from a Chinese patient who had handled and eaten pork before admission to hospital (44). We found immunization with 2015033 protected mice from the fatal challenge of these different STs in CC1; to our knowledge, this is the first live vaccine reported to provide STs-independent immunity against *S. suis*.

In summary, our results described immunization with a five-gene deletion attenuated mutant 2015033, which simultaneously expressed CPS and various immunogenic proteins, suppressed lethal STSLS, and conferred cross-STs immunity in mice during *S. suis* infection. Taken together, these results indicate that multi-gene deletion may be a promising strategy for the development of vaccines against encapsulated invasive pathogens.

## MATERIALS AND METHODS

### Bacterial strains, plasmids and growth conditions

Bacterial strains and plasmids used in this study are listed in Table S1. Primers used to construct gene-deleting plasmids are listed in Table S2. Primers used to identify the gene deletion mutant are listed in Table S3. *S. suis* strains were cultured using Tryptic Soy Broth (Becton, Dickinson) or Tryptic Soy Agar (Becton, Dickinson) plus 10% newborn bovine serum at 37 °C. *E. coli* DH5α was cultured in LB broth or on LB agar at 37 °C. Spectinomycin (Sigma) was supplemented at 50 μg/μL for *E. coli* and 100 μg/μL for *S. suis* when required.

### Cell cultures

Cells used in this study are listed in Table S1. Caco-2 were cultured in DMEM with 10% fetal bovine serum (FBS), 1% L-glutamine and 1% nonessential amino acids. A549 were cultured in F-12K medium with 10% FBS. HBMEC were established by Professor Kwang Sik (Johns Hopkins University School of Medicine) and shared by Dr. Xiangru Wang (Huazhong Agricultural University) (45). HBMEC were cultured in RPMI1640 medium supplemented with 10% FBS, 1% sodium pyruvate, 2% L-glutamine, 1% vitamins, 2% essential amino acids, 1% nonessential amino acids, and 1% penicillin and streptomycin. All cells were incubated at 37°C with 5% CO_2_.

### Construction of the five-gene (*sly*,*scpA*, *ssnA*, *italic>fhb*, and *ssads*) deletion mutant

Target genes are listed in Table S4 and were deleted by an allelic replacement strategy as previously described (37). Five recombinant plasmids pSET4s-*sly*, pSET4s-*scpA*, pSET4s-*ssnA*, pSET4s-*fhb* and pSET4s-*ssads* were generated. pSET4s-*sly* was subsequently introduced into *S. suis* SC19 by electroporation. The *sly* gene deletion mutant Δ*sly* was selected due to its sensitivity to spectinomycin. The pSET4s-*scpA* was then introduced into the Δ*sly* mutant to generate a double-gene deletion mutant and additional deletions of *ssnA*, *fhb* and *ssads* were performed sequentially following the protocol described above to generate a five-gene deletion mutant.

### Biological characteristics of 2015033

The procedure of sample preparation for the transmission electron microscopy was described previously (46). Morphology was observed with an FEI GI 20 TWIN transmission electron microscope (Wuhan, China) at an accelerating voltage of 200 kV. Antibiotic susceptibility of SC19 or 2015033 to penicillin was determined using an *E*-test according to manufacturer’s instructions (Biomerieux).

### Human blood bactericidal assay

Blood assays were conducted according to an approval issued by the Medical Ethics Committee of the Huazhong Agricultural University (Wuhan, China). The bactericidal assay of *S. suis* strains in human blood was performed as described previously (46). Briefly, diluted cultures of SC19 or 2015033 (100 μL, 1 × 10^3^ CFU) was mixed with fresh heparinized human venous blood (900 μL), and the mixtures were rotated moderately at 37 °C. Samples were taken at 1, 2 and 3 h, and the viable bacteria were determined by plating diluted samples on TSA plates. The growth index was defined as the ratio of increased bacterial counts after incubation over the original bacterial counts [(CFU_Tn_-CFU_T0_) / CFU_T0_].

### Hemolytic activity assay

SC19 or 2015033 (5 × 10^7^ CFU, 100 μL) were incubated with 4% sheep erythrocytes (900 μL) for 1 h at 37 °C. Tryptic Soy Broth with 10% newborn bovine serum was used as a negative control. Subsequently, 200 μL supernatant fluids were transferred to a 96-well plate and were measured at 540 nm with a microplate reader (Bio-Tek, synergy, HT).

### Cell experiments

Cytotoxicity assays were conducted using lactate dehydrogenase (LDH) cytotoxicity detection kits (Beyotime) following the recommendation of the manufacturer. Briefly, 1 × 10^4^ cells were infected with 2 × 10^5^ CFU of SC19 or 2015033 and incubated for 4 h at 37 °C with 5% CO_2_. After incubation, the supernatant was collected and the absorbance was read at 490 nm using a microplate reader (Bio-Tek, synergy, HT). The percentage of cytotoxicity was calculated using the formula: Cytotoxicity (%) = (LDH release from infected cells – spontaneous release of LDH from uninfected cells) / (maximum LDH release from cell lysate – spontaneous release of LDH from uninfected cells) × 100%. To identify the death model of HBMEC after *S. suis* infection, 2 × 10^6^ cells were harvested 8 h after treatment with SC19 or 2015033 (MOI = 20), washed in PBS and stained by PI/Annexin V-FITC double stain Kit (Beyotime). The samples were immediately analyzed by flow cytometry (FACSVerse, BD Biosciences). To perform the cell adherence assays, HBMEC cells were grown to confluence as a monolayer in 24-well plates averaging 1 × 10^5^ cells per well. HBMEC cells were incubated with *S. suis* (1 × 10^6^ CFU) for 3 h. After washing three times with PBS, the cells were lysed using 1% saponin and bacterial adhesion was determined by plating the lysates on TSA plates. Percentage of adherence bacteria was expressed as (CFU _on plate_ / CFU _in original inoculum_) × 100%.

### Experimental infections of mice

Six-week-old female specific-pathogen-free (SPF) BALB/C mice were used in this study. Animal experiments were approved by Committee on the Ethics of Animal Experiments, Huazhong Agricultural University (Wuhan, China). The survival study was performed as follows: mice (ten mice per group) were intraperitoneally infected with 2 × 10^8^ CFU of SC19 or 2015033 (suspended in 200 μL PBS). Survival time was monitored and recorded for 2 weeks post infection. Experimental mice for bacterial load were challenged intraperitoneally with *S. suis* strains (non-lethal dose of 5 × 10^7^ CFU, suspended in 200 μL PBS), bacteria counts in blood and tissue samples (lung, liver, and brain) were quantified at 24 h and 72 h post-infection.

### Immunization and challenge

Four-week-old female SPF BALB/c mice (n = 114) were divided into two groups. Mice in one group were subcutaneously immunized with 2 × 10^7^ CFU of 2015033 (suspended in 200 μL PBS), while mice in the other group were subcutaneously injected with 200 μL PBS, which served as the negative control group. The immunization was performed twice at a 2-week interval. Mice in the two groups were intraperitoneally infected with ST7 strain SC19, ST1 strain P1/7, or ST658 strain LSM102 (2× 10^8^ CFU, suspended in 200 μL PBS, ten mice per subgroup). Mice were then daily monitored up to 14 days post-infection to calculate survival rate.

### Opsonic killing assay

Blood samples were collected on day 14 after the booster immunization. Sera were separated by centrifugation (3000 × g, 5 min). ST7 strain SC19, ST1 strain P1/7, or ST658 strain LSM102 (1 × 10^3^ CFU) was suspended in 100 μL immune or nonimmune mouse sera and incubated for 30 min at 37 °C. The mixtures were then added into 400 μL fresh nonimmune mouse whole blood for 1 h incubation with gentle rotation at 37 °C. Diluted samples were plated on TSA plates to determine bacterial counts. Survival percentages of *S. suis* strains incubated with nonimmune sera were regarded as 100% and of *S. suis* strains incubated with immune sera were regarded as the corresponding relative value.

### Quantification of Ab responses

Blood samples were collected on days 7, 14, 21, and 28 after immunization. Sera were separated by centrifugation (3000 × g, 5 min). To quantify Ab responses specific to SC19, polyvinylchloride 96-well plates were coated with 1 × 10^7^ CFU of SC19 strain for 24 h at 37 °C and then washed with PBST. The washed plates were blocked with 5% skim milk for 2 h at 37 °C. Mouse sera were diluted in 5% skim milk and incubated for 1 h at 37 °C. Plates were then washed with PBST and incubated with peroxidase-conjugated goat anti-mouse IgG, IgG1, and IgG2a (Southern Biotech). Plates were then coated with 3,3’,5,5’-Tetramethylbenzidine (TMB) substrate and 2M H_2_SO_4_ was used to stop the reaction. Absorbance was read at 450 nm. For quantification of Ab responses specific to the proteins of *S. suis*, polyvinylchloride 96-well plates were coated with a nonencapsulated mutant Δ*cps*2E and other experimental operations were performed following the protocol described above.

### Quantification of mouse cytokines

To evaluate the inflammation-inducing ability of 2015033, six-week-old female mice (n = 3) were inoculated intraperitoneally with SC19 or 2015033, respectively (5 × 10^7^ CFU, suspended in 200 μL PBS). At 18 h post infection, blood samples were harvested by cardiac puncture. Sera were separated by centrifugation (3000 × g, 5 min). Levels of IL-6, IL-10, IL-12p70, IFN-γ, TNF-α, and MCP-1 were measured by the cytometric bead array (CBA) mouse inflammation kit (BD Biosciences) according to the manufacturer’s instructions. To evaluate the STSLS-suppressing ability of the immunization, on day 28, immunized or non-immunized mice (n = 3) were intraperitoneally infected with SC19 (5 × 10^7^ CFU, suspended in 200 μL PBS) or PBS only. At 24 h post-infection, blood samples were harvested by cardiac puncture. Sera were separated by centrifugation (3000 × g, 5 min). Other experimental operations were performed following the protocol described above.

### Population snapshot of clonal complexes

Data of 1003 different STs were obtained from the *S. suis* MLST website (available at https://pubmlst.org/) sited at the University of Oxford. Population snapshot of clonal complexes was drawn using eBURST software (10).

### Statistical analysis

All statistical analyses were performed using GraphPad Prism version 5.0. Data were expressed as the mean ± SD. Unpaired, two-tailed Student’s *t*-test was used to analyze the statistical significance between two groups. Two-way ANOVA with Tukey’s multiple-comparison test was used to analyze the statistical significance among more than two groups. The log-rank test was used to analyze the statistical significance of survival curves.

## ACKNOWLEDGMENTS

We thank Ying Zhou and Liangliang Gao for their technical support. We thank Dr. Sekizaki (National Institute of Animal Health, Japan) for sharing plasmid pSET4s and Xiangru Wang for supplying HBMEC.

The study was supported by the National Key R&D Program of China (2017YFC1600105), the National Natural Science Foundation of China (31502080, 31772083), the Fundamental Research Funds for the Central Universities (2662017JC040, 2662016QD010) and the Open Project Program of Jiangsu Key Laboratory of Zoonosis (R1605).

All authors declare no conflicts of interest.

---

**Fig S1.**
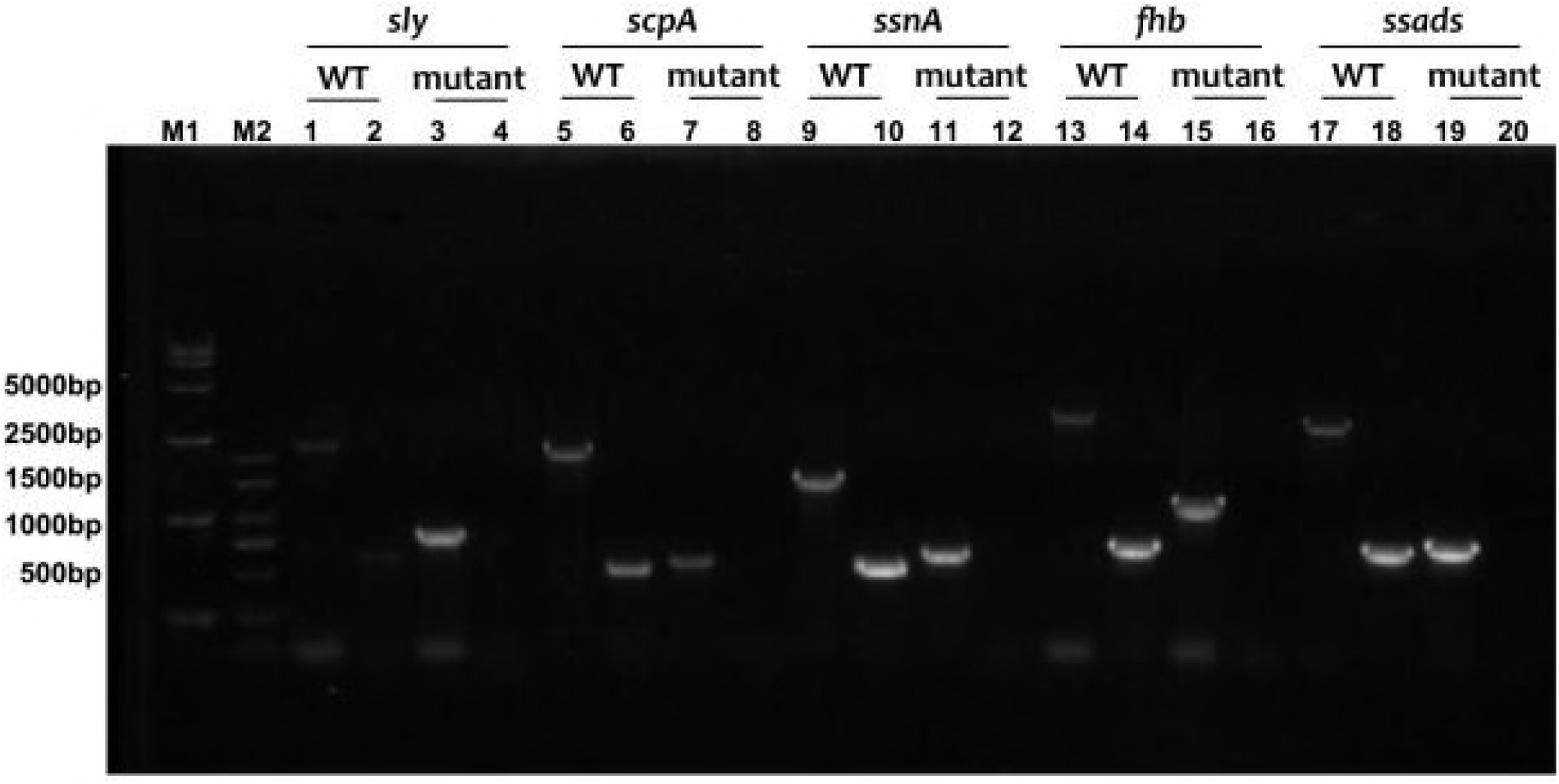
Confirmation of target deletion by PCR. Successful construction of the five-gene (*sly*, *scpA*, *ssnA*, *fhb*, and *ssads*) deletion mutant was determined by PCR using primers listed in Table S3.

**Table S1** Information of strains and plasmids used in this study.

**Table S2** Primers used to amplify the flanking sequence of target genes.

**Table S3** Primers used to identify the gene deletion mutant.

**Table S4** Information of target genes.

